# Evolution of miR-155 and its probable targets: *MST1R, Adam10*, and *CD9* genes expression levels in adults and children gastritis patients with *H. pylori* infection

**DOI:** 10.1101/2023.04.07.536047

**Authors:** Ramina Mahboobi, Fatemeh Fallah, Abbas Yadegar, Roshanak Shams, Amir Sadeghi, Naghi Dara, Mojdeh Hakemi-Vala

## Abstract

**Introduction:** *H. pylori* accounts for the main factor of gastric cancer which contributes to an immune response with the promotion of inflammation. Recently, the effect of miRNAs on the prognosis of diseases is gaining the attraction of investigators. Herein, we studied the expression levels of miR-155 and its identified targets (*MST1R, Adam10*, and *CD9*) along with miR-155 expression correlation with virulence factors of *H. pylori* (*vacA* and *cag genes*), *H. pylori* colonization, and inflammation, in patients’ candidates for gastric endoscopy due to gastritis.

**Methods:** in this study total of 50 and 26 biopsy samples were taken from adults and children respectively. Biochemical and molecular identification of samples was performed using culture and PCR of *ureC, 16sRNA*, along with amplification of *vacA* and *cagA* genes for pathogenicity of bacteria. The qRT-PCR was carried out using STEM-LOOP RT-PCR (dye-based) for the evaluation of miR-155 expression level and *Adam10, CD9*, and *MST1R* expression levels. All Real-Time PCR reactions were carried out in triplicate and data analysis was conducted using REST.

**Results:** A total of 30 out of 17 biopsy samples in adults and children were positive for *H. pylori* in both PCR and culture, respectively. The expression level of miR-155 is closely related to the *H. pylori infection* and the down-regulation of *CD9*, and *MST1R* genes in *H. pylori* (+) samples compared to *H. pylori* (-) in adults’ biopsy (p=0.0001). Although, there wasn’t any relation between *cagA* and *vacA* genes with the expression of miR-155 in evaluated biopsy samples in both adults and children.^i^

**Conclusion:** This study for the first time revealed that the expression of *MST1R, CD9*, and *adam10 genes* was relatively related to the expression of miR-155, and indicated that the miR-155 overexpression promoted the poor prognosis of *H. pylori* infection in adults.

## Introduction

*Helicobacter pylori* (*H. pylori*) - a gram-negative spiral-shaped bacterium - has infected about 50% of the worldwide population (1) with a prevalence range of 85-95% and 30-50% in developing and developed countries, respectively (2, 3). More than 4.4 billion people are infected with *H. pylori* worldwide, so although the sanitation and level of hygiene are improved in recent years, the prevalence of infection is still the highest, especially in developing countries (4). The bacteria are colonized in gastric mucosa and mostly are promoted asymptomatic, although in about 20% they developed the gastrointestinal disease such as acute or chronic gastritis and peptic ulcer. *H. pylori* infection even can cause gastric cancer which accounts for the fifth most common cancer with third high mortality rates in each gender (5), and according to the World Health Organization(WHO), *H. pylori* is classified as the first carcinogen that caused gastric cancer (6).

*H. pylori* colonization contributes to an immune response with the promotion of inflammation, however, the mechanism of inflammation induction following infection with *H. pylori* is not fully understood (7). Recently, the effect of micro-RNAs (miRNAs) - non-protein-coding RNAs of ∼22 nucleotides - in the prognosis of diseases has been gained the attraction of investigators and various studies had been demonstrated the role of miRNAs on the pathogenesis of *H* .*pylori* infection(8-10). It has been observed that miRNAs would render the inflammation and immune induction in response to infection (11). Furthermore, miRNAs could alter the host’s response to infection and would limit the replication and distribution of infectious microorganisms (8).

The studies have demonstrated that miR-155 is proficiently involved in the up-regulation of pro-inflammatory response in *H. pylori* infection(12). The miR-155 plays a pivotal role in the induction of innate immune responses mediated by macrophages, lymphocytes, and dendritic cells. This miRNA is up-regulated in response to toll-like receptor (TLR) ligands or TNF-α expression which are dependent on NF-kB and AP-1 transcription factors (13, 14). It has been shown that the expression of miR-155 would negatively reduce the pathogenicity of *H. pylori*, so, the miR-155 expression accounts for a potential factor indicating the pathogenesis of *H. pylori* infection (15). *H. pylori* promotes the secretion and translocation of pathogenic proteins like *CagA* and *VacA* to down regulate the signaling pathways such as NF-kB, therefore it contributes to the induction of miR-155 expression through the NF-kB signaling pathway (15). Hence, nowadays, besides evaluation of common virulence factors of *H. pylori* such as *vacA* and *cagA*, genes investigation of miRNAs expression profile would be helpful to anticipate the phenomenon of disease.

In our previous study, using bioinformatics analysis we predicted the target genes related to miR-155 and we found *MST1R, Adam10*, and *CD9* genes as three top targets which were related to *H. pylori* and the occurrence of gastric cancer (under publish). Herein, we studied the expression levels of miR-155 and it’s identified targets (*MST1R, Adam10*, and *CD9*) along with their correlation with virulence factors of *H. pylori* (*vacA* and *cagA genes*), *H. pylori* colonization, and inflammation, in patients’ candidates for gastric endoscopy due to gastritis. In this research, the biopsy specimens of patients with gastric diseases related to *H. pylori* in adults and children were used. We supposed that the study on co-expression of miR-155 and its targets as well as *H. pylori* virulence factors would render the prognosis of metaplasia and gastric cancer.

## Materials and methods

### Study population

#### Patients

A total of 50 adult patients (20–70-year-old) who were referred to the Research Center for Gastroenterology and Liver Diseases in Taleghani hospital (Tehran, Iran) along with 26 patients (1–12-year-old) who were referred to the Mofid children’s hospital (Tehran, Iran) during 2021-2022, were enrolled in this research. Patients were diagnosed with the symptoms of dyspeptic and gastritis by the Gastrologist and didn’t receive antibiotics 4 weeks before endoscopy. All patients have signed the informed consent form before the endoscopy. This study was approved by the ethical community of Shahid Beheshti University of medical science with the number IR.SMBU.MSP.REC.1399.756.

#### Samples

For each case, four biopsy samples were obtained from the mucosa of the antrum. One biopsy sample was transferred to the Thioglycolate media for culture. The second biopsy sample was transferred to the Brucella broth media and kept at -80°C for DNA extraction. The third biopsy sample was fixed at formalin and sent to the pathologist for pathological evaluation; And the fourth biopsy was transferred to the RNA FIX solution, kept at 4°C for 24h initially, and then transferred to the -20°C refrigerator for RNA extraction.

### Biochemical and Molecular testing of *H. pylori* for identification of strain and genotype

#### Culture and biochemical testing

About 76 biopsy samples in Thioglycolate media were transferred to the BHI (Brain Heart Infusion) broth supplemented with 10% bovine serum in an aseptic condition. Then the tissue was grinded and homogenized followed by transferring to the Brucella Agar composed of 7% sheep blood, 10% FCS (fetal cow serum), and the selective campylobacter antibiotics (vancomycin 2mg, polymyxin B 0.05 mg, Trimethoprim 1 mg, and amphotericin b 2.5 mg). they were cultured at microaerophilic incubator adjusted with 10% CO_2_, 5% O_2_, and 85% N_2_ and evaluated on the third, fifth, and seventh day of culture. The confirmation of *H. pylori* was carried out using biochemical tests including, catalase and urease tests following gram staining on cultured bacteria.

#### DNA extraction

A tissue Genomic DNA extraction mini kit (Favorprep, Pingtung-Taiwan) was used for DNA extraction according to the manufactured structure. Briefly, for isolation of DNA from tissue: biopsies were ground using cell lysis and proteinase K solution, then the lysed solution was bound to the membrane using the binding buffer and washed twice using wash buffer and eluted using elution buffer. For DNA extraction from cultured cells, initially, the colonies were dissolved in culture solution and centrifuged, then the supernatant was discarded and followed according to DNA extraction of tissue samples. DNA samples were kept at -20°C before further evaluation.

#### Molecular confirmation

To confirm the type and strains of bacteria, the presence of 16s rRNA, *ureC*, and *glmM* genes were determined using specific primer sets and for the pathogenicity of *H. pylori*, the presence of *vacA* and *cagA* genes were also considered using specific primers. The list of all primers is indicated in Table1. The PCR program was carried out as follows: initial denaturation at 94°C for 4 min, following 30 cycles of denaturation at 94°C for 1 min, annealing at 58°C, 56°C, 52°C, and °C 57 for 45sec, for *16s rRNA, glmM, cagA, and vacA* receptively, and extension at 72°C for 1 min, with a final extension at 72°C for 15sec. The PCR products were electrophoresis using 8% agarose gel and visualized under UV light.

**Table 1.**
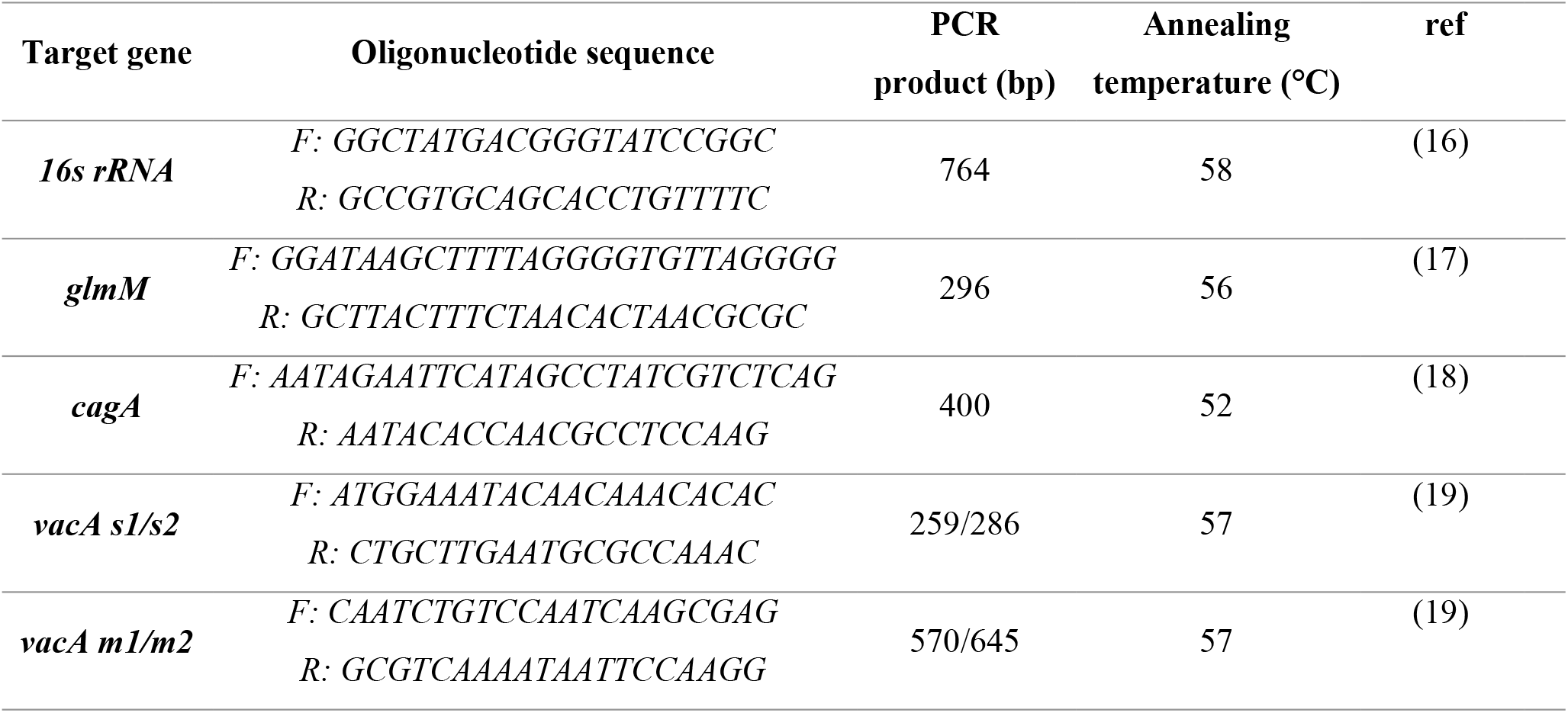
List of primers used for PCR

### Evaluation of the expression levels of miR-155 and *MST1R, Adam10*, and *CD9* probable target genes using qRT-PCR

#### RNA isolation and cDNA synthesis

Total RNA was isolated using TRIzonl solution (Biobasic Inc, Canada) according to the manufactured structure and DNase was added to remove any DNA from the extracted samples. Because, the miRNAs are small RNA with 22 nucleotides, therefore to increase the length of the RNA template, the STEM-LOOP RT specific primers of RNU6: *CTCAACTGGTGTCGTGGAGTCGGCAATTCAGTTGA* and miR-155: *CTCAAGGTGTCGTGGAGTCCCCAATTCAGTTGAGACCCTAT* were designed in which the length of RNA exceed up to >60 nucleotides. Then, the cDNA synthesis was performed immediately with STEM-LOOP RT specific primers and incubated at 80°C for 5 min following incubation at 50°C for another 5 min at thermocycler. At the next step, the cDNA synthesis was followed using an Easy cDNA synthesis kit (Parstous Inc, Mashhad, Iran) according to the manufactured structure and initially incubated at 50°C for 60 min then incubated at 75°C for 15 min at a thermocycler. cDNA samples were kept at -80°C until further evaluation.

#### Quantitative Real-Time PCR for evaluation of miR-155 expression level

The qRT-PCR was carried out using STEM-LOOP RT-PCR (dye-based) method in which the forward primers added more nucleotides to the target cDNA and optimized the melting temperature of the RT-PCR reaction while the universal revered primers were applied to break down the pin-shaped structure of cDNAs; as well, the *RNU6* housekeeping gene was applied as an internal control. Detection of miR-155 and *RNU6* genes were performed according to the followed reaction: A total volume of 20 μl reaction with 10 μl of master mix, 0.5 μl of each primer, 2 μl cDNA were prepared and then placed at RT-thermocycler with the following program: initial denaturation at 94°C for 4 min, following 30 cycles of denaturation at 94°C for 1 min, annealing at 58°C for 45sec, and extension at 72°C for 1 min, with a final extension at 72°C for 15sec.

### Cyber Green Real-time PCR for evaluation of probable target genes expression levels

Evaluation of expression levels of miR-155 target genes (*Adam10, CD9*, and *MST1R*) was carried out by the Cyber Green Real-time PCR method. A volume of 10μl of Cyber Green master mix, 0.5 μl of each forward and reverse primers, and 2 μl of cDNA with a total volume adjusted to 20 μl using deionized water. The *β-actin* gene was applied as an internal control and the RT-PCR program was carried out for 45 cycles with activation at 95°C for 15sec, denaturation at 95°C for 20sec, and annealing at 57°C for 40sec.

Specific primers for miR-155 and *Adam10, CD9*, and *MST1R* genes for Real-time PCR reactions are listed in Table 2.

**Table 2.**
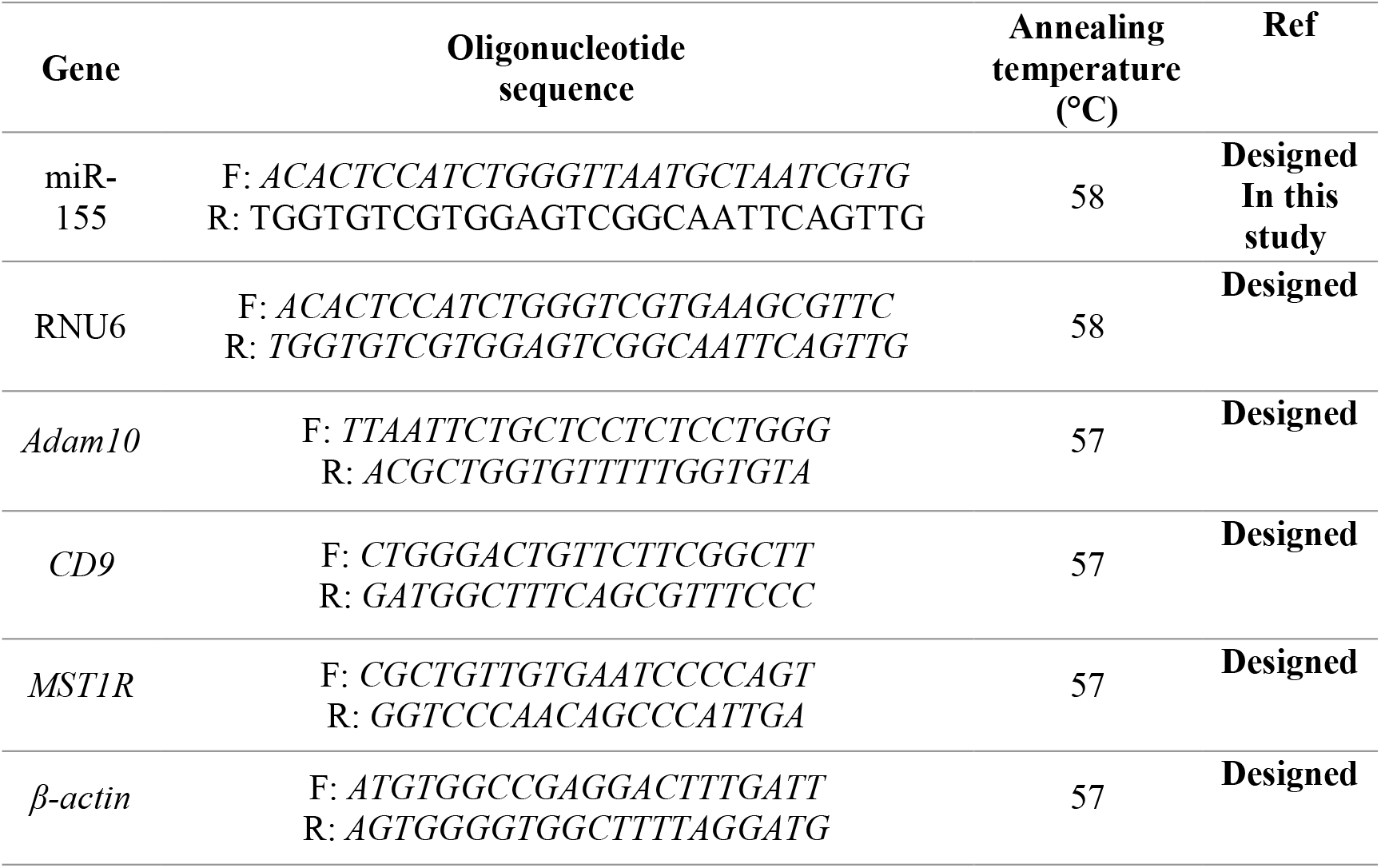
List of specific primers used for qRT-PCR

### Statistical analysis

All Real-Time PCR reactions were carried out in triplicate and data analysis was conducted using REST (Relative Expression Software Tool) 2009 for Ct values. All statistical analysis was performed using SPSS v.20 (SPSS incl, Chigaco, IL., USA) and GraphPad Prism 6. The numerical data were presented as mean ± standard deviation. The independent t-test and One-way ANOVA test were applied for evaluating the expression of genes in normal and patients’ biopsy samples. A *p*-value of less than 0.05 was considered significant statistically.

## Results

### The presence of *H. pylori* in biopsy samples was confirmed by culture, biochemical and molecular tests

#### Culture and Biochemical tests

The biopsy samples were cultured at Brucella agar for three days and among 50 biopsy samples obtained from adults, 30 biopsy samples were positive and the grey colonies were observed at culture media. Also, 5 biopsy samples out of 26 biopsy samples were obtained from children were positive for *H. pylori* in culture media. The Catalase and urease tests of all positive cultures were positive. The *H*.*pylori* colonization and inflammation grad were reported according to Sydney Histopathological grading system (20) The results are represented in Table 3.

**Table 3.**
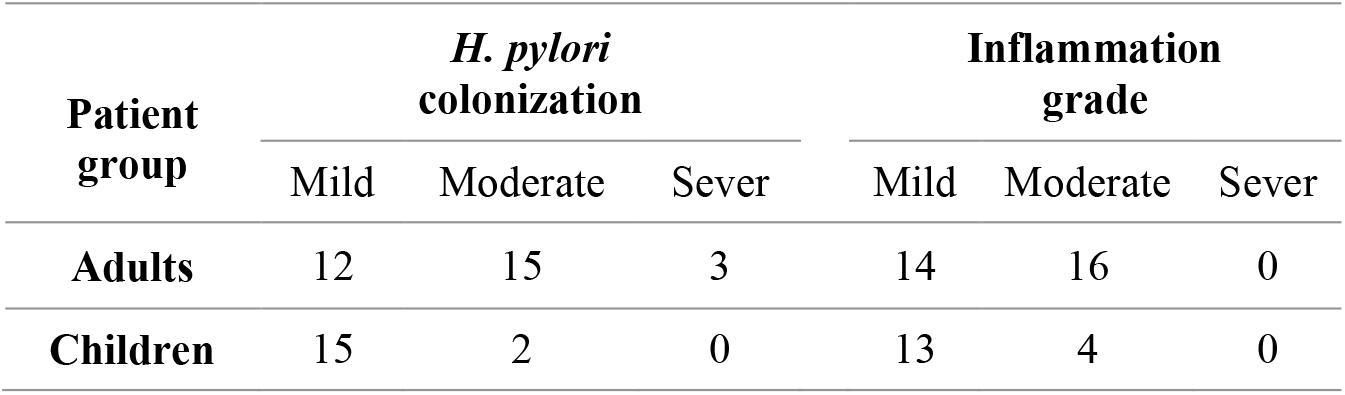
*H*.*pylori* colonization and inflammation grade in biopsy of gastritis patients

| Patient group | *H. pylori* colonization |  |  | Inflammation grade |  |  |
| --- | --- | --- | --- | --- | --- | --- |
|  | Mild | Moderate | Sever | Mild | Moderate | Sever |
| Adults | 12 | 15 | 3 | 14 | 16 | 0 |
| Children | 15 | 2 | 0 | 13 | 4 | 0 |

#### Molecular study for H. pylori genotyping

A total of 30 out of 50 positive *H. pylori* in culture media obtained from adults were positive for the presence of *16S rRNA* and *ureC* genes in PCR evaluation. In biopsy samples obtained from children, 12 samples in addition to 5 positive cultured samples were positive for the presence of *ureC* and *16S rRNA* genes in PCR evaluation. The prevalence of the *cagA* gene in adults with positive *H. pylori* was 63.34%, whereas, among 17 *H. pylori* positive samples obtained from children, 70.5% were positive for the presence of the *cagA* gene. The results are represented in Table 4 and Figure 1.

**Table 4.**
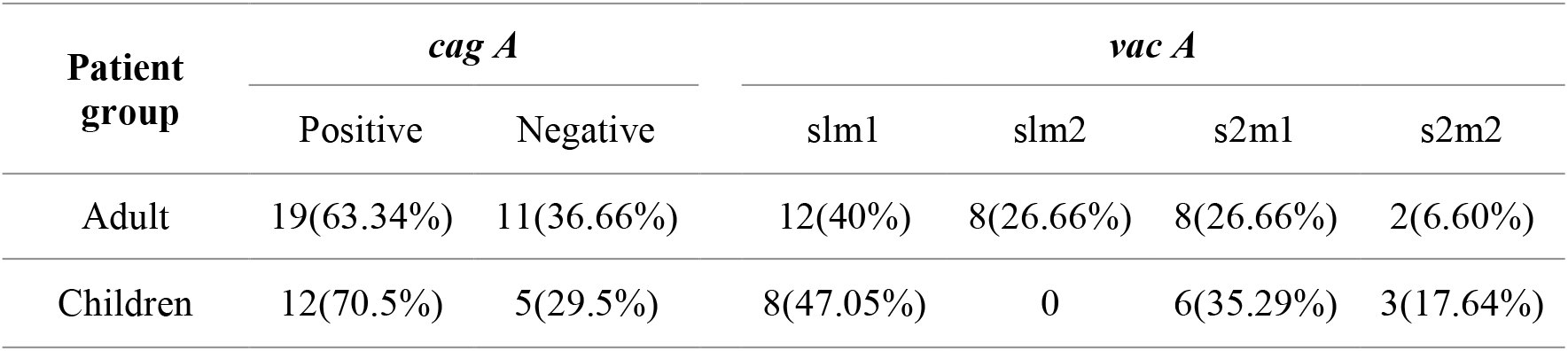
The number of patients samples infected by *H. pylori* with and without *cagA* and *vacA* virulence factor

**Figure 1.**
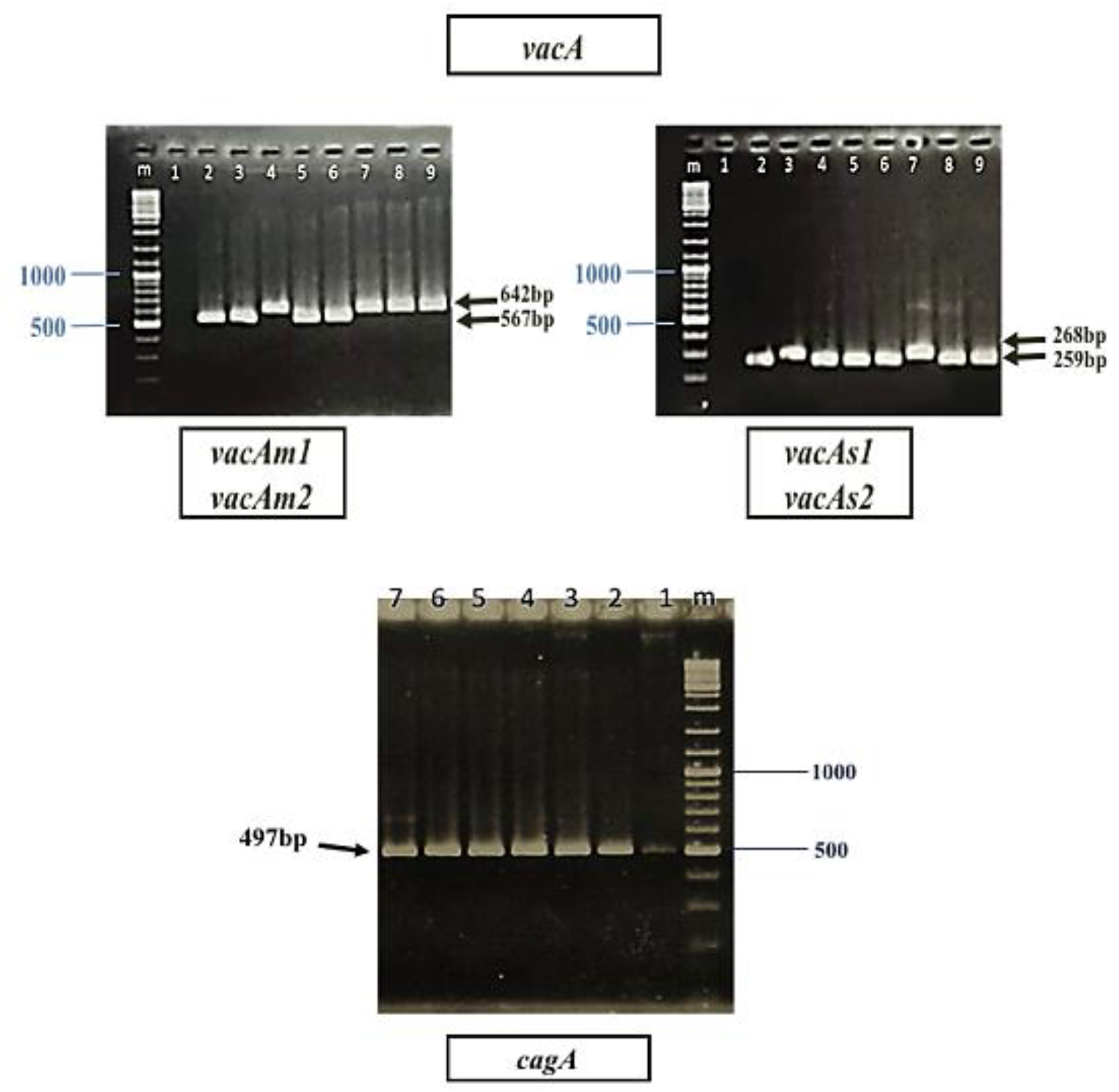
Genotyping of the different alleles of *vacA* and *CagA* gene in gel electrophoresis. A) the bad in size of 567 bp represents a *vacA m1* allele (lanes 2,3,5, and 6), and the band in size 642 bp represent a VacA *m2* allele (lanes 4,7, 6, and 9), lane 1 is a negative control, lane M 1Kb DNA ladder. B) the bad in size of 259 bp represents a *vacA s1* allele (lanes 4,5,6, and 9), and the band in size 286 bp represent a *Va*cA *s2* allele (lanes 2,3, and 7), lane 1 is a negative control, lane M 1Kb DNA ladder. C) the PCR of *cagA* gene with 497bp in gentle electrophoresis (lane 2-7), lane 1 is negative control, lane M 1Kb DNA ladder

### Comparsion of the expression level of miR-155 in *H*.*pylori* (+) samples in adults versus children

The qRT-PCR analysis showed the high expression level of miR-155 in *H. pylori* (+) compared to *H. pylori* (-) biopsy samples in adults, indicating the statistically significant relation between the high expression level of miR-155 in the occurrence of *H. pylori* in adults (p=0.0001). While the high expression level of miR-155 in *H. pylori* (+) in comparison with *H. pylori* (-) biopsy samples in children was not statistically related to *H. pylori* infection (*p*=0.190). The results are represented in Table 4 and Figure 2.

**Table 4.** The miR-155 expression levels in adult and children gastritis biopsy samples

| Patient group | *H. pylori* (+) | *H. pylori* (-) | Mean difference | *p*-value |
| --- | --- | --- | --- | --- |
| miR-155 (Adults) | 25.21±6.28 | 15.10±3.52 | 10.11 | 0.0001 |
| miR-155 (Children) | 18.61±4.79 | 14.76±7.60 | 3.85 | 0.190 |

**Figure 2.**
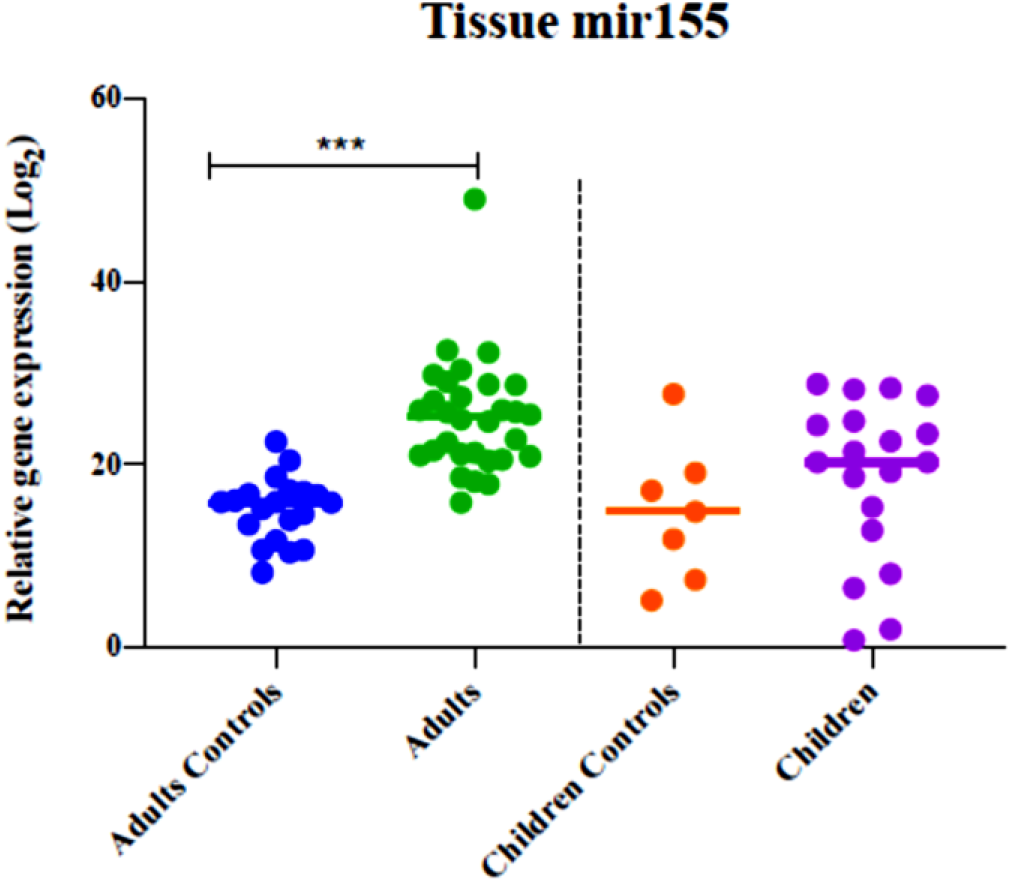
Expression levels of miR-155 in adults (*H. pylori* (+)) and children (*H. pylori* (+)) compared to controls *(H. pylori* (-) gastritis patients.

### The expression level of *Adam10, CD9*, and *MST1R* showed differences in *H. pylori* (+) and (-) samples

The different expression levels of *Adam10, CD9*, and *MST1R* genes were observed in *H. pylori* (+) and (-) samples. Up-regulation of the *Adam10* gene and down-regulation of *CD9*, and *MST1R* **genes** in *H. pylori* (+) samples compared to *H. pylori* (-) was statistically significant in adults’ biopsies (p=0.0001); demonstrating the effect of *H. pylori* infection on *Adam10, Cd9*, and *MST1R* expression levels in adult patients. In contrast, up-regulation of *Adam10, CD9*, and *MST1R* in *H. pylori* (+) compared to *H. pylori* (-) samples was not statistically significant (*p*=0.67, 0.5, 0.67 for *Adam10, CD9*, and *MST1R*, respectively) in children’s biopsies. All results were indicated in Table 5 and Figure 3.

**Table 5.** Target genes expression levels in adults and children biopsy samples

| Target genes |  | *H. pylori* (+) | *H. pylori* (-) | *p*-value | Mean difference between adults and children |
| --- | --- | --- | --- | --- | --- |
| *Adam10* | Adults | 8.25±2.96 | 16.22±7.14 | 0.0001 | 6.60 |
|  | Children | 4.60±6.67 | 3.58±4.12 | 0.67 |  |
| *CD9* | Adults | 8.58±1.66 | 13.18±9.66 | 0.0001 | 9.08 |
|  | Children | 6.21±7.01 | 4.09±7.21 | 0.5 |  |
| *MST1R* | Adults | 1.88±6.38 | 18.33±5.6 | 0.0001 | 11.70 |
|  | children | 6.62±4.80 | 6.33±5.86 | 0.67 |  |

**Figure 3.**
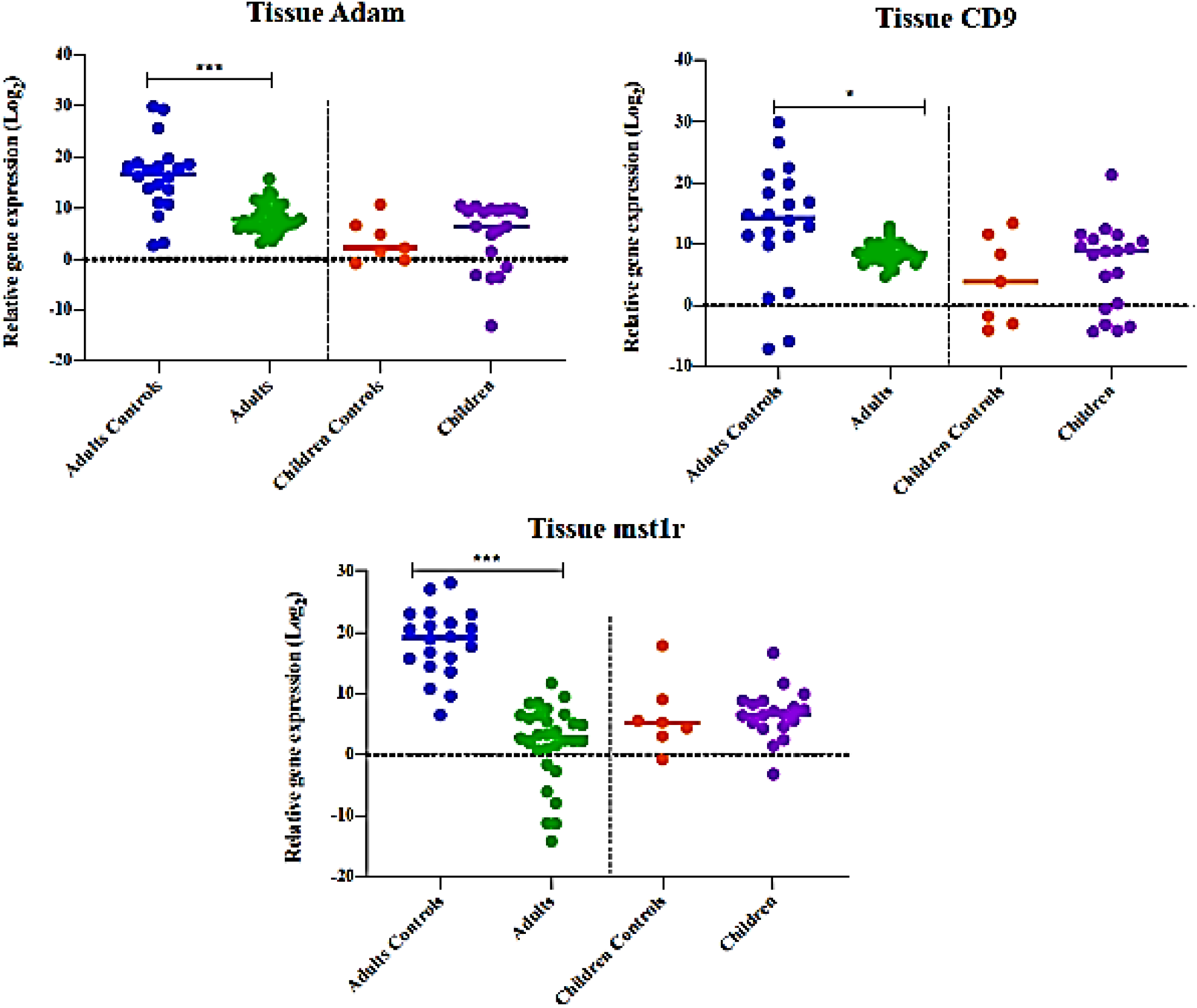
Expression levels of *Adam10, Cd9*, and *MST1R* in adults (*H*.*pylori* (+)) and children (*H*.*pylori* (+)) compared to controls (*H*.*pylori* (-)) gastritis patients

### Correlation between *cagA* and *vacA* virulence genes with the expression of miR-155 in gastritis patients

In adult biopsy samples, down-regulation of the miR-155 was observed in *cagA* and *vacA genes* positive samples in comparison to negative samples. While the miR-155 in *cagA* and *vacA genes* positive children biopsy samples up-regulated. However, the miR-155 expression level in *cagA* positive samples in either adults or children compared with *cagA* gene negative samples indicated that *cagA gene* didn’t correlate with the expression of miR-155 in either group (p=0.2). Also, there wasn’t any significant correlation between the presence of the *vacA* gene and the expression of miR-155 in adults and children as well. The results were indicated in Table 6.

**Table 6.**
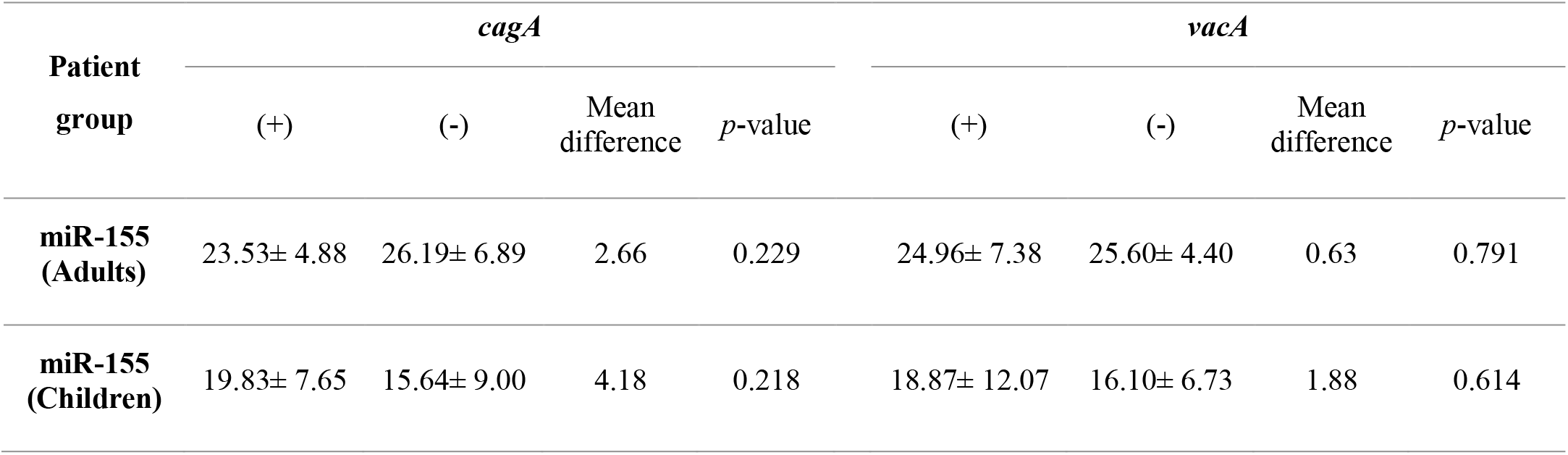
The expression level of miR-155 in *cagA* and *vacA* positive and negative biopsy samples

### Correlation between *H. pylori* colonization and inflammation grade with the expression of miR-155

The colonization level of *H. pylori* and grade of inflammation directly related to the higher expression of miR-155 in adults but the colonization and inflammation didn’t statistically relate to the expression of miR-155 in children. The results are indicated in Table 7.

**Table 7.**
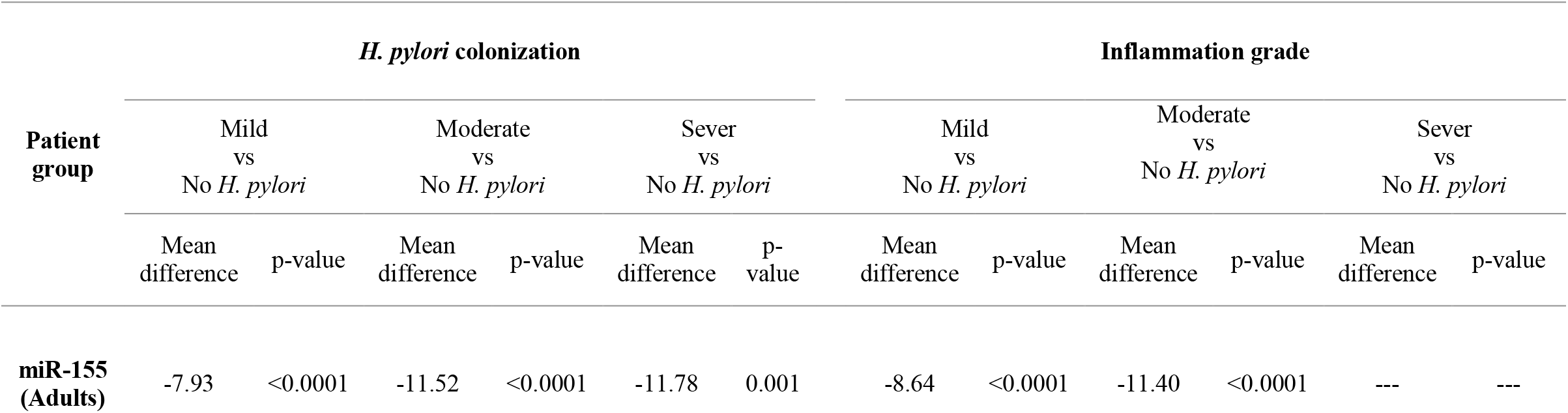
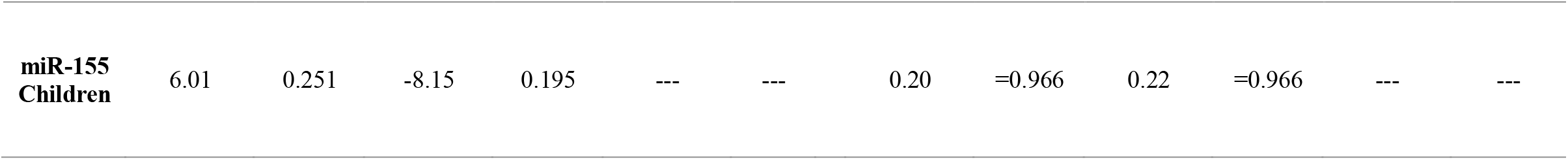
Correlation between miR-155 expression level and *H. pylori* colonization and inflammation grade

| Patient group | *H. pylori* colonization |  |  |  |  |  | Inflammation grade |  |  |  |  |  |
| --- | --- | --- | --- | --- | --- | --- | --- | --- | --- | --- | --- | --- |
|  | Mild vs No *H. pylori* |  | Moderate vs No *H. pylori* |  | Sever vs No *H. pylori* |  | Mild vs No *H. pylori* |  | Moderate vs No *H. pylori* |  | Sever vs No *H. pylori* |  |
|  | Mean difference | *p*-value | Mean difference | *p*-value | Mean difference | *p*-value | Mean difference | *p*-value | Mean difference | *p*-value | Mean difference | *p*-value |
| **miR-155 (Adults)** | -7.93 | <0.0001 | -11.52 | <0.0001 | -11.78 | 0.001 | -8.64 | <0.0001 | -11.40 | <0.0001 | --- | --- |
| miR-155<br>Children | 6.01 | 0.251 | -8.15 | 0.195 | --- | --- | 0.20 | =0.966 | 0.22 | =0.966 | --- | --- |

## Discussion

The gut microenvironment is sterile until the baby gives birth and *H. pylori* is acquired during childhood without a specific indication, due to the immunologic tolerance of *H. pylori* in childhood, the bacteria persist in the human. Therefore, *H. pylori* is considered a human pathogen in later years and offer potential drawback at a younger age. Decreased eradication rates and increased drug resistance are the main problems in children (21). The clinical manifestation of *H. pylori* infection is gastritis, peptic ulcer, atrophy, and gastric cancer which is classified as a major cause of gastric cancer according to National Institution of Health (NIH) NIH. Recently, various studies have been indicated the role of miRNAs in the development of clinical manifestation of *H. pylori* in the gastrointestinal tract (13). This current study aimed to investigate the expression level of miR-155 in adults and children who were candidates for gastric endoscopy and find the relation between miR-155 with inflammation, and *H. pylori* virulence factors as well as *H. pylori* colonization, and the expression of miR-155 target genes.

Initially, Matsushima and colleagues characterized the expression of miRNAs in gastric mucosa of patients with *H. pylori* using high throughput analysis, they discovered that 31 miRNAs were expressed in the gastric mucosa (14). Later, Santos et. al demonstrated that the expression of miR-150-5p, miR-155-5p, and miR-3163 is altered during *H. pylori* infection, they confirmed that the miR-155 overexpression accounts for the main target for the prevention of cancer, especially in the early stage of cancer (15). In our study, 30 samples out of 50 samples obtained from adults and 17 samples out of 26 samples obtained from children were positive for *H. pylori* infection either based on culture or molecular confirmations (Table 3). As well, the presence and frequency of the *cagA* and *vacA* virulence factor genes were confirmed in adults and children (Table 4, Figure 1). According to qRT-PCR analysis, indicating the high expression level of miR-155 in *H. pylori* (+) compared to *H. pylori* (-) biopsy samples in adults, we found the statistically significant relation between the high expression level of miR-155 in the occurrence of *H. pylori* in adults (p=0.0001). While the high expression level of miR-155 was not statistically related to *H. pylori* infection (p=0.190) in *H. pylori* (+) compared to *H. pylori* (-) biopsy samples in children (Table 4 and Figure 2). Our results from adults patients were in accordance with the studies that demonstrated the proficiency involvement of miR-155 up-regulation in pro-inflammatory response during *H. pylori* infection (22, 23), Also, one study revealed that the expression of miR-155 would negatively reduce the pathogenicity of *H. pylori*, so, the miR-155 expression accounts as a potential factor indicating the pathogenesis of *H. pylori* infection (15).

We investigated the correlation between miR-155 expression level and its three top probable target genes *(Adam10, CD9*, and *MST1R*) and observed no significant down-regulation of *Adam10* (*p*=0.67*), CD9 (p=*0.5), and *MST1R* (*p*=0.67) in *H. pylori* (+) compared to *H. pylori* (-) samples in children biopsy. Whereas, significant down-regulation of *Adam10* gene, *CD9*, and *MST1R* genes in *H. pylori* (+) samples compared to *H. pylori* (-) was observed in adults’ biopsy (p=0.0001) (Table 5 and Figure 3).

*Adam10 gene*, is a part of the Adam family overexpressed in malignancies and cancer development (24). Yuan-Yu Wang et al also revealed that the *Adam10* was associated with a poor prognosis of gastric cancer and induced the progression and metastasis of cancer (25). Previous studies indicated that *CD9* gene expression decreased in oral squamous cell carcinoma and increased in gastric cancer. The molecular mechanism of *CD9* is apoptotic signaling while ligated to the gastric cells and it is over-expressed in gastric cancer (26). *MST1R gene*, is a member of the tyrosine kinase family which is potentially associated with epithelial cancer such as gastric cancer; the expression of miR-155-5p contributes to the higher expression of *MST1R* and severe pathogenicity of *H. pylori* in adults but there wasn’t any study considering the effect of *MST1R* expression on the expression of miR-155. Therefore, for the first time, the finding of the current study revealed that the expression of the *MST1R* gene is related to the severe phenomenon of gastric disease and negatively related to the expression level of miR-155 (27). Overall, our results declared a positive correlation between miR-155 expression with *Adam10* as well as a negative correlation between miR-155 expression with *CD9* and *MST1R* expression in adult patients.

As mentioned earlier, *H. pylori* promotes the secretion and translocation of pathogenic proteins like *cagA* and *vacA* to down regulate the signaling pathways such as NF-kB, consequently contributing to the induction of miR-155 expression through the NF-kB signaling pathway. Cheng et.al declared that the miR-155 over-expressed in patients with *cagA* positive *H. pylori (28)*. But we couldn’t find any statistical correlation between *cagA* and *vacA* genes and the expression of miR-155 (Table 6). However, we suppose that the miR-155 expression renders the inflammation and colonization of *H. pylori* in gastric mucosa.

The prevalence of *H. pylori* depends on age and geographical distribution that the prevalence of *H. pylori* in children less than 10 years old was reported lower than in adults aged over 20 whereas, in developing countries, the rate of infection is higher in childhood (21). We observed that the incidence of *H. pylori* was higher in adults that reason was due to the initial colonization of bacteria in childhood and immune tolerance of the body. We also focused on the correlation between miR-155 expression level and *H. pylori* colonization and inflammation grade and additionally found that the expression of miR-155 was statistically correlated with higher *H. pylori* colonization and inflammation in adults but the colonization and inflammation didn’t statistically relate to the expression of miR-155 in children (Table 7).

## Conclusion

This current study aimed to study the expression levels of miR-155 and its identified targets (*MST1R, Adam10*, and *CD9*) in the biopsy specimens of patients’ candidates for gastric endoscopy due to gastritis related to *H. pylori* in adults and children. This study for the first time indicated that the expression of *MST1R, CD9*, and *Adam10* was closely related to the expression of miR-155, although the *MST1R* had a statistical correlation with miR-155 expression and indicated that the miR-155 overexpression promoted the poor prognosis of *H. pylori* infection in adults. We also evaluated the miR-155 expression and its correlation with virulence factors of *H. pylori* (*vacA* and *cagA genes*), *H. pylori* colonization, and inflammation. Our study couldn’t find any statistical correlation between *cagA* and *vacA* virulence factors and the expression of miR-155 but we found that the expression of miR-155 was statistically correlated with higher *H. pylori* colonization and inflammation in adults. Finally, according to our results obtained from evaluation on biopsy samples of adults and children, we supposed that the study on co-expression of miR-155 and its target genes as well as *H. pylori* virulence factors and colonization would render the prognosis of metaplasia and gastric cancer in adult patients. These findings introduced the new therapeutic factors to scheme targeted gene therapy of gastric cancer along with early detection of the severe form of the disease to prevent the progression of gastric cancer.

